# Robust alignment of single-cell and spatial transcriptomes with CytoSPACE

**DOI:** 10.1101/2022.05.20.488356

**Authors:** Milad R. Vahid, Erin L. Brown, Chloé B. Steen, Minji Kang, Andrew J. Gentles, Aaron M. Newman

## Abstract

Recent studies have emphasized the importance of single-cell spatial biology, yet available assays for spatial transcriptomics have limited gene recovery or low spatial resolution. Here we introduce CytoSPACE, a method for aligning single-cell and spatial transcriptomes via convex linear optimization. Across diverse platforms and tissue types, we show that CytoSPACE outperforms previous methods with respect to noise-tolerance, accuracy, and efficiency, enabling improved analysis of spatial transcriptomics data at single-cell resolution.

## Main

Single-cell spatial organization is a key determinant of cell state and function. For example, in human tumors, local signaling networks differentially impact individual cells and their surrounding microenvironments, with implications for tumor growth, progression, and response to therapy^1-6^. While spatial transcriptomics (ST) has become a powerful tool for delineating spatial gene expression in primary tissue specimens, commonly used platforms, such as 10x Visium, remain limited to bulk gene expression measurements, where each spatially-resolved expression profile is derived from as many as 10 cells or more^7^.

Accordingly, a number of computational methods have been developed to infer cellular composition in a given bulk ST sample^8-23^. Most such methods use reference profiles derived from single-cell RNA sequencing (scRNA-seq) data to deconvolve ST spots into a matrix of cell type proportions. However, these methods lack single-cell resolution, hindering the discovery of spatially-defined cell states, their interaction patterns, and their surrounding communities (**Supplementary Fig. 1**).

To address this challenge, we developed cellular (Cyto) Spatial Positioning Analysis via Constrained Expression alignment (CytoSPACE), an efficient computational approach for mapping individual cells from a reference scRNA-seq atlas to precise spatial locations in a bulk ST dataset (**Fig. 1a, Supplementary Fig. 1**). Unlike related methods^24,25^, we formulate single-cell/spot assignment as a convex optimization problem and solve this problem using a novel application of the Jonker-Volgenant shortest augmenting path algorithm^26^. Our approach guarantees an optimal mapping result while exhibiting improved noise tolerance (**Methods**). The output is a reconstructed tissue specimen with both high gene coverage and spatially-resolved scRNA-seq data suitable for downstream analysis, including the discovery of context-dependent cell states. On both simulated and real ST datasets, we find that CytoSPACE substantially outperforms related methods for resolving single-cell spatial composition.

**Figure 1:**
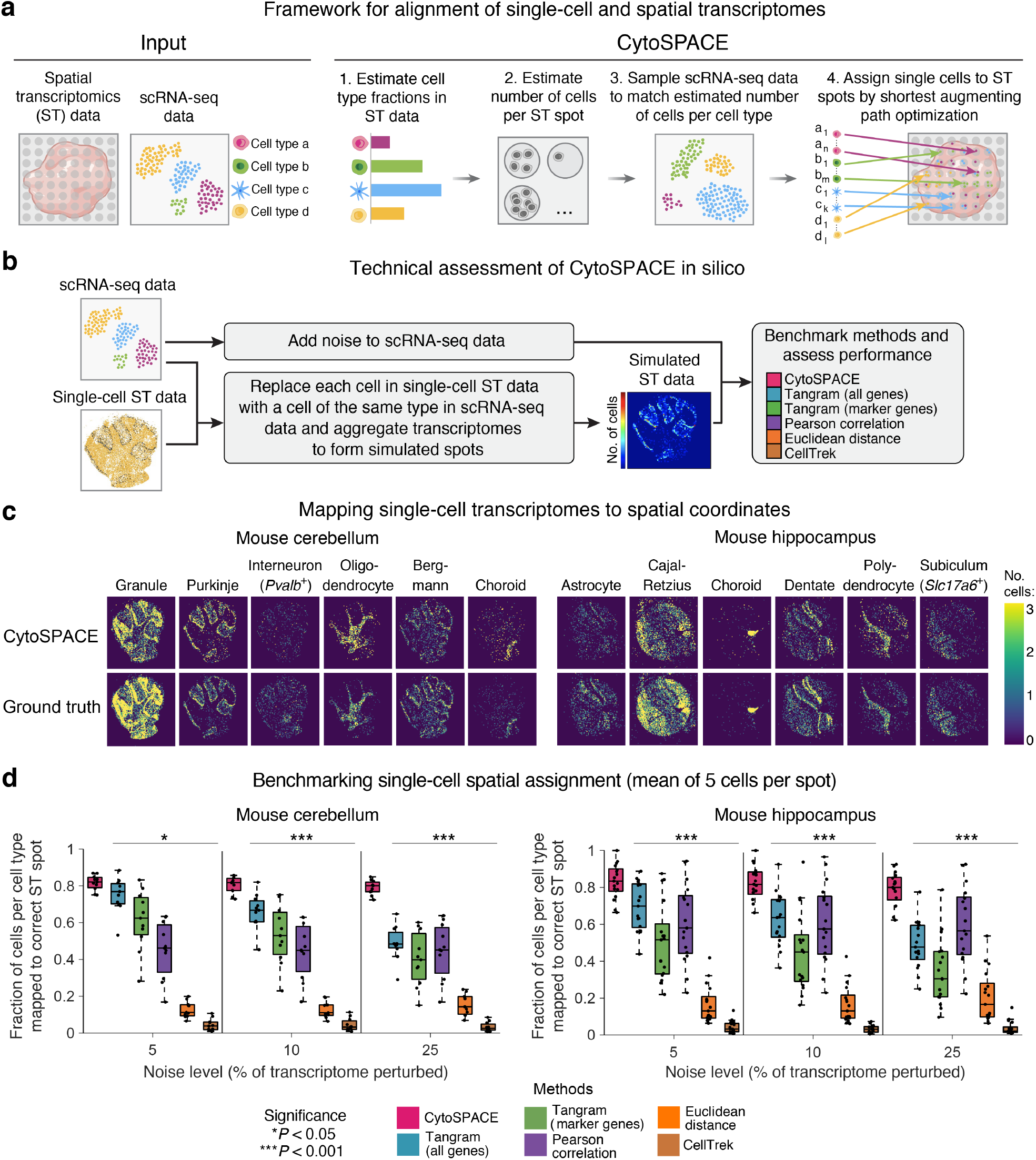
Development and technical assessment of CytoSPACE. **a**, Schematic of a typical CytoSPACE workflow. Given an ST dataset *A* and an annotated scRNA-seq dataset *B* where the latter covers major cell types in *A*, CytoSPACE consists of the following key steps: (1) application of an existing ST deconvolution method (e.g., Spatial Seurat, RCTD) to estimate cell type fractions in *A* using reference profiles from *B*, (2) estimation of the number of cells per spot in *A*, (3) sampling of *B* to match the inferred number of cells per cell type in *A*, and (4) alignment of single-cell and spatial transcriptomes (*A* → *B*) using shortest augmenting path-based optimization. **b–d**, Technical assessment of CytoSPACE. The labels *a*_1_, *a*_*n*_, …, *d*_1_, *d*_*n*_ denote individual single cells of cell type *a, a*, …, *d, d*, respectively. **b**, Framework for evaluating CytoSPACE using simulated ST datasets with fully-defined single-cell composition and spot resolution (**Methods**). **c**, Heat maps depicting CytoSPACE performance for aligning scRNA-seq data (with 5% added noise) to spatial locations in ST datasets simulated with 5 cells per spot, on average (**Methods**). Only cell types with distinct spatial structure are shown here for clarity. For all evaluated cell types, see **Supplementary Figure 4. d**, Performance across distinct methods, mouse brain regions, and noise levels for assigning individual cells to the correct spot in simulated ST datasets (**Methods**). Each point represents a single cell type (mouse cerebellum, *n* = 11; mouse hippocampus, *n* = 17). The box center lines, box bounds, and whiskers indicate the medians, first and third quartiles and minimum and maximum values within 1.5× the interquartile range of the box limits, respectively. Statistical significance was assessed relative to CytoSPACE using a two-sided paired Wilcoxon test and reported as the maximum *P* value per noise level (\**P* < 0.05, \*\*\**P* < 0.001).

CytoSPACE proceeds in four main steps (**Fig. 1a**). First, to account for the disparity between scRNA-seq and ST data in the number of cells per cell type, two parameters are required: (i) the fractional abundance of each cell type within the ST sample and (ii) the number of cells per spot. The former is determined using an external deconvolution tool, such as Spatial Seurat^14^, RCTD^18^, SPOTlight^20^, cell2location^27^, or CIBERSORTx^28^. By default, the latter is directly inferred by CytoSPACE using an approach for estimating RNA abundance (**Methods**). Once both parameters are estimated, the scRNA-seq dataset is randomly sampled to match the predicted number of cells per cell type in the ST dataset. Up-sampling is done for cell types with insufficient representation, either by drawing with replacement or by introducing placeholder cells (**Methods**). Finally, CytoSPACE assigns each cell to spatial coordinates in a manner that minimizes a correlation-based cost function constrained by the inferred number of cells per spot via a shortest augmenting path optimization algorithm. An efficient integer programming approximation method that yields comparable results is also provided^29^ (**Methods**).

To assess the performance of CytoSPACE, we began by simulating ST datasets with fully defined single-cell composition. For this purpose, we leveraged previously published mouse cerebellum (*n* = 11 major cell types) and hippocampus (*n* = 17 major cell types) data generated using Slide-seq, a platform with high spatial resolution (approximately single cell) but limited gene coverage^30^ (**Fig. 1b**). To increase transcriptome representation, we first replaced each Slide-seq bead with a single-cell expression profile of the same cell type derived from an scRNA-seq atlas of the same brain region^31^ (**Methods**). We then superimposed a spatial grid with tunable dimensions in order to pool single-cell transcriptomes into pseudo-bulk transcriptomes. This was done across a range of realistic spot resolutions (mean of 5, 15, and 30 cells per spot). To guarantee a unique spatial address for every cell in the scRNA-seq query dataset, we created a paired scRNA-seq atlas from the cells underlying each pseudo-bulk ST array. Finally, to emulate technical and platform-specific variation between scRNA-seq and ST datasets, we added noise in varying amounts to the scRNA-seq data (**Supplementary Fig. 2**; **Methods**).

Next, we evaluated methods for CytoSPACE parameter inference. For cell type enumeration, we employed Spatial Seurat, which showed strong concordance with known proportions in simulated ST datasets (**Supplementary Fig. 3a**). To approximate the number of cells per spot, we implemented a simple approach based on RNA abundance estimation, which was significantly correlated with ground truth expectations in simulated ST samples (**Supplementary Fig. 3b, Methods**).

We then benchmarked CytoSPACE against two recently described algorithms for scRNA-seq and ST alignment: Tangram, which integrates scRNA-seq and ST data via maximization of a spatial correlation function using nonconvex optimization^24^; and CellTrek, which uses Spatial Seurat^14^ to identify a shared embedding between scRNA-seq and ST data and then applies random forest modeling to predict spatial coordinates^25^. We also assessed two naïve approaches, Pearson correlation and Euclidean distance. To compare outputs, each cell was assigned to the spot with the highest score (all approaches but CellTrek) or the spot with the closest Euclidean distance to the cell’s predicted spatial location (CellTrek only). Further details are provided in Methods.

Remarkably, across multiple evaluated noise levels and cell types, CytoSPACE achieved significantly higher precision than other methods for mapping single cells to their known locations in simulated ST datasets (**Fig. 1c,d; Supplementary Figs. 4 and 5**). This was true for multiple spatial resolutions independent of brain region, both for individual cell types and across all evaluable cells (**Fig. 1d and Supplementary Fig. 5**). We also observed similar results with an independent method for determining cell type abundance in ST data (RCTD^18^), demonstrating robustness (**Supplementary Fig. 6**). These data highlight the potential of CytoSPACE to deliver improved spatial mapping of scRNA-seq data.

To evaluate performance on real ST datasets, we next examined primary tumor specimens from three types of solid malignancy: melanoma, breast cancer, and colon cancer. In total, six scRNA-seq/ST combinations, encompassing six bulk ST samples (*n* = 4 Visium; *n* = 2 legacy ST), including one HER2+ formalin fixed paraffin embedded (FFPE) breast tumor specimen and three scRNA-seq datasets from matching tumor subtypes, were analyzed^32-35^. All cell types in each scRNA-seq dataset were aligned by CytoSPACE (**Fig. 2a and Supplementary Fig. 7**) and compared to Tangram and CellTrek (**Supplementary Fig. 7**). Notably, CytoSPACE was substantially more efficient than other methods, processing a Visium-scale dataset in approximately 5 minutes on a single CPU core (**Supplementary Fig. 8a**). This was true regardless of whether we applied shortest augmenting path or integer programming approximation approaches, both of which achieved comparable results (**Supplementary Fig. 8b**). To quantitatively compare the recovery of cell states with respect to spatial localization patterns in the tumor microenvironment (TME), we dichotomized assigned cells into two groups within each cell type by their proximity to tumor cells. We then assessed whether gene sets marking TME cell states with known localization were skewed in the expected orientation (**Fig. 2b**; **Methods**).

**Figure 2:**
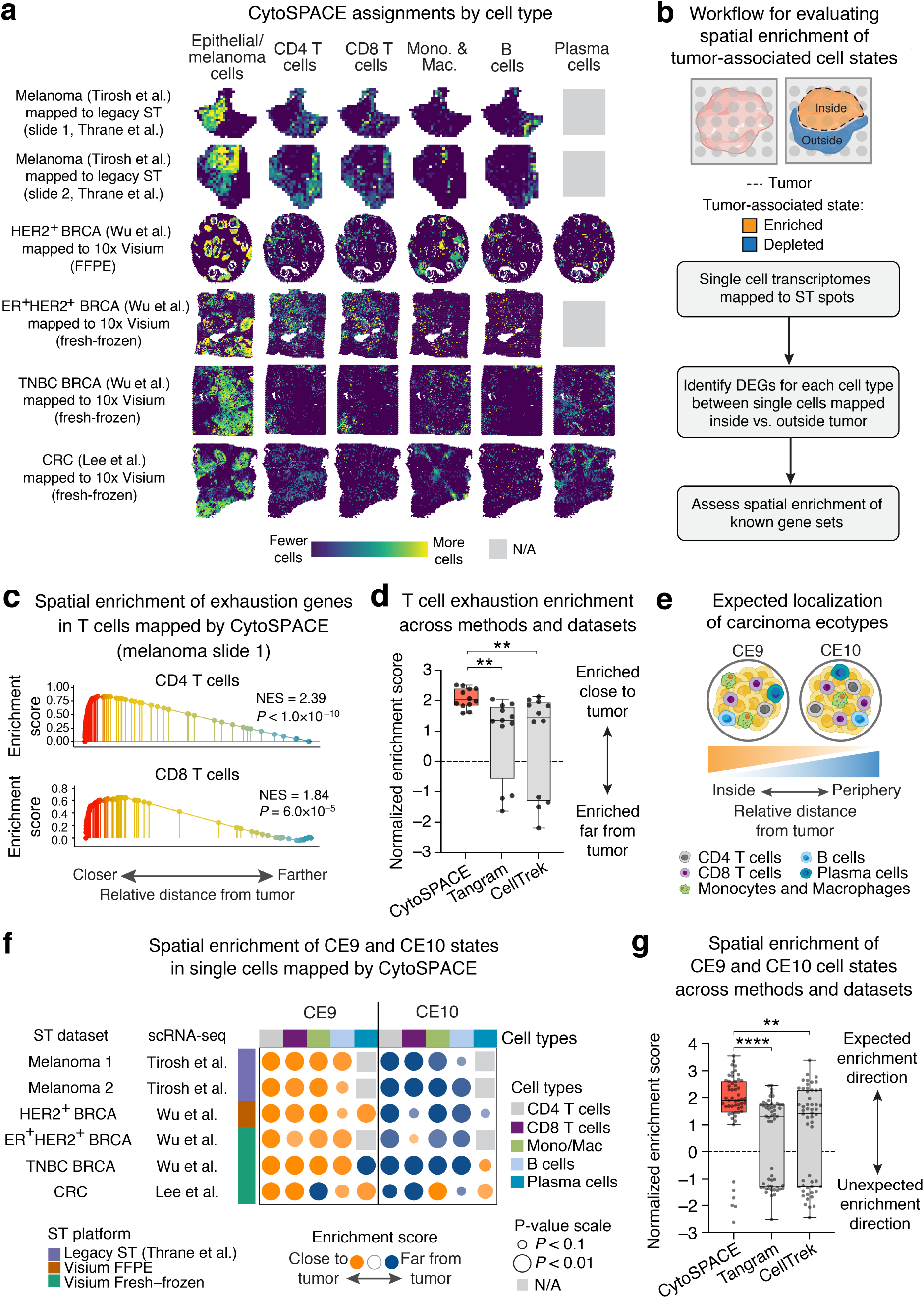
Single-cell cartography of TME communities. **a**, Heat maps showing diverse scRNA-seq tumor atlases mapped onto clinically-matched ST target datasets by CytoSPACE. Details of scRNA-seq and ST datasets are provided within the first and second set of parentheses, respectively. Only cell types analyzed in subsequent panels are shown (for all mapped cell types, see **Supplementary Fig. 7**). N/A, missing from author-supplied scRNA-seq annotations; BRCA, breast cancer; CRC, colorectal cancer; FFPE, formalin-fixed paraffin embedded. **b**–**g**, Identification, validation, and benchmarking analysis of tumor-associated cell states. **b**, Procedure for evaluating whether cell-type-specific states are preferentially localized to the tumor core or periphery. Differentially expressed genes (DEGs) were identified between cells stratified into two groups based on their distance from tumor cells (**Methods**) and assessed for spatial enrichment by pre-ranked gene set enrichment analysis (GSEA). **c**–**d**, Evaluation of T cell exhaustion. **c**, Spatial enrichment of T cell exhaustion genes in melanoma-associated CD4 and CD8 T cell transcriptomes mapped by CytoSPACE to a melanoma specimen profiled by ST (slide 1, Thrane et al.). NES, normalized enrichment score. **d**, Same as c but showing CD4 and CD8 T cell enrichment scores for 6 evaluated scRNA-seq/ST pairs (*n* = 12 values per box) across 3 methods. **e**, Schematic of TME communities with expected localization to the tumor core (CE9) and periphery (CE10) in carcinomas and melanoma. **f**, Bubble plot showing whether CE9 and CE10-specific cell states are enriched within the tumor core (orange) or periphery (blue) in single cells mapped to ST spots by CytoSPACE. The NES and statistical significance of pre-ranked GSEA are denoted by bubble color and size, respectively. Single-cell RNA-seq datasets without annotated plasma cells are indicated by gray boxes. **g**, Same as f but comparing NES across 3 methods (*n* = 54 values per box). To unify the expected enrichment direction of CE9 and CE10 cell states, normalized enrichment scores for the latter were multiplied by –1. In d and g, the box center lines, box bounds, and whiskers denote the medians, first and third quartiles and minimum and maximum values, respectively. Group comparisons in d and g were determined relative to CytoSPACE via a two-sided, paired Wilcoxon test (\*\**P* < 0.01, \*\*\*\**P* < 0.0001).

We started by considering T cell exhaustion, a canonical state of dysfunction arising from prolonged antigen exposure in tumor-infiltrating T cells^36^. Consistent with expectation, CytoSPACE recovered spatial enrichment of T cell exhaustion genes^37^ in CD4 and CD8 T cells mapped closest to cancer cells in all six scRNA-seq and ST dataset combinations (**Fig. 2c,d; Supplementary Fig. 9a**). In contrast, Tangram and CellTrek produced single cell mappings with significantly lower enrichment of T cell exhaustion genes in the expected orientation, with 25% to 33% of cases showing enrichment in the opposite direction, away from the tumor core (**Fig. 2d; Supplementary Fig. 9a**).

To demonstrate applicability to other spatially-biased cell states, we next extended our analysis to diverse TME lineages, identifying cell type-specific genes that vary in expression as a function of distance from tumor cells. To validate our results, we considered two recently defined cellular ecosystem subtypes in human carcinoma, CE9 and CE10^4^. These “ecotypes,” which were also observed in melanoma, each encompass B cells, plasma cells, CD8 T cells, CD4 T cells, and monocytes/macrophages with stereotypical spatial localization. CE9 cell states are preferentially localized to the tumor core whereas CE10 states are preferentially localized to the tumor periphery^4^ (**Fig. 2e**). Using marker genes specific to each state^4^, we asked whether single cells mapped by each method were consistent with CE9 and CE10-specific patterns of spatial localization (**Fig. 2e**). Indeed, as observed for T cell exhaustion factors, CytoSPACE successfully recovered expected spatial biases in CE9 and CE10 cell states across lymphoid and myeloid lineages (**Fig. 2f**), significantly outperforming Tangram and CellTrek in both the magnitude and orientation of marker gene enrichments (**Fig. 2g; Supplementary Fig. 9b,c**). As further validation, we analyzed predicted spatial localization patterns of *TREM2*+ and *FOLR2*+ macrophages, which were recently shown to localize to the tumor stroma and to the tumor mass, respectively, across diverse cancer types^6^ (**Supplementary Fig. 10a**). Only CytoSPACE recapitulated these prior findings with statistical significance (**Supplementary Fig. 10b**). Moreover, when inferred spatial locations were projected onto scRNA-seq data, single cells generally failed to organize as a function of distance from tumor cells (**Supplementary Fig. 11**). These data underscore the ability of CytoSPACE to accurately identify spatially-resolved cell states, including those not discernible from scRNA-seq data alone.

In summary, CytoSPACE is a new tool for aligning single-cell and spatial transcriptomes via convex linear optimization. Unlike related methods, CytoSPACE ensures a globally optimal single-cell/spot alignment conditioned on a correlation-based cost function and the number of cells per spot. Moreover, it can be readily extended to accommodate additional constraints, such as the fractional composition of each cell type per spot (e.g., as inferred by RCTD^18^ or cell2location^27^). In contrast, CellTrek is dependent on the co-embedding learned by Spatial Seurat, which can erase subtle, yet important biological signal (e.g., cell state differences), as was recently shown^38^. While Tangram is robust in idealized settings, it cannot guarantee a globally optimal solution. While CytoSPACE requires two input parameters, both parameters can be reasonably well-estimated using standard approaches, suggesting they are unlikely to pose a major barrier in practice. Furthermore, on both simulated and real datasets, CytoSPACE was substantially more accurate and efficient than related methods. As such, we anticipate that CytoSPACE will prove immediately useful for deciphering single-cell spatial variation and community structure in diverse physiological and pathological settings.

## Acknowledgements

We thank A. Chaudhuri, W. Zhang, and M. Matusiak for providing critical feedback on this manuscript. We are grateful to A. Bergersen for assistance with software development and N. Midler, N. Semenkovich, W. Zhang, and M. Matusiak for assistance with software testing. This work was supported by grants from the National Cancer Institute (A.M.N., R01CA255450 and R00CA187192; A.J.G., R21CA238971 and U54CA209971), the Virginia and D.K. Ludwig Fund for Cancer Research (A.M.N.), the Stinehart-Reed Foundation (A.M.N.), the Stanford Bio-X Interdisciplinary Initiatives Seed Grants Program (IIP) (A.M.N.), and the Donald E. and Delia B. Baxter Foundation (A.M.N.).

## Author Contributions

M.R.V. and A.M.N. conceived of the study; M.R.V., E.L.B., C.B.S., and A.M.N. wrote the paper; and M.R.V., E.L.B., C.B.S., and A.M.N. performed data analysis and interpretation with assistance from A.J.G. M.R.V., E.L.B., and C.B.S. developed and implemented CytoSPACE and prepared documentation with assistance from A.M.N. M.K. assisted with data curation. All authors commented on the manuscript at all stages.

## Methods

### CytoSPACE analytical framework

CytoSPACE leverages convex linear optimization to efficiently assign single-cell transcriptomes to spatial transcriptomics data. To formulate the assignment problem mapping individual cells in scRNA-seq data to spatial coordinates in ST data, let an *N* × *C* matrix *A* denote single-cell gene expression profiles with *N* genes and *C* cells; let an *M* × *S* matrix *B* denote gene expression profiles of spatial transcriptomics (ST) data with *M* genes and *S* spots; and let *G* be the vector of length *g* that contains the subset of desired genes shared by both data sets. For both gene expression profile matrices, values are first normalized to counts per million (or transcripts per million for platforms covering the full gene body) and then transferred into log_2_ space. Next, we estimate the number *n*_s_, *s* = 1, …, *S*, of cells contributing RNA content in the *s*^*th*^ spot of ST data (see “Estimating the number of cells per spot”). We assume that the *s*^*th*^ spot contains *n*_s_ sub-spots that can each be assigned to a single cell, and build an *M* × *L* matrix 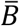 by replicating the *s*^*th*^ column of *B, n*_s_ times, where 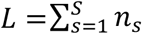 denotes the total number of estimated sub-spots in the ST data. As described in the following sections, we then sample the scRNA-seq matrix *A* such that the total number of cells, with cell types represented according to their inferred fractional abundances, matches the total number of columns in 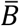, yielding an *N* × *K* matrix *Ā*, where *K* = *L*. Define an assignment *x* := [*x*_*kl*_], *k* = 1, …, *K* and *l* = 1, …, *L*, such that *x*_*kl*_ = 1 if the *k*^*th*^ cell in the scRNA-seq data is assigned to the *l*^*th*^ sub-spot in the ST data, and *x*_*kl*_ = 0 otherwise. We find the optimal cell/sub-spot assignment *x*^*^ that minimizes the following linear cost function:

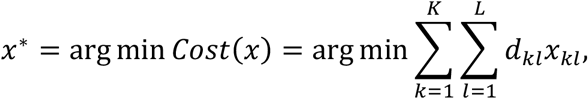

subject to:

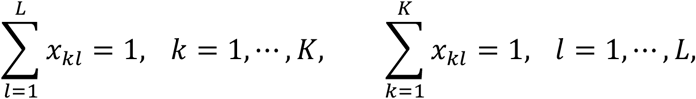

where *d*_*kl*_ denotes the distance between the gene expression profiles of the *k*^*th*^ cell and the *l*^*th*^ sub-spot. The above constraints guarantee that each cell is only assigned to one sub-spot and each sub-spot only receives one cell. In general, *d*_*kl*_ can be obtained using any metric that quantifies the similarity between the gene expression profiles of the reference and target data sets. We examined different similarity metrics for simulated data and selected Pearson correlation as below due to its robustness to noise:

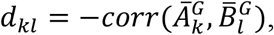

where 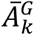 and 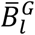 denote the *k*^*th*^ and *l*^*th*^ columns of expression matrices *Ā* and 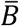, respectively, for the shared genes in *G*.

We provide two possible solvers for CytoSPACE, both of which will return the globally optimal solution of the above problem as formulated. The first of these implements the shortest augmenting paths-based Jonker-Volgenant algorithm, in which we solve the dual problem of the above formulation defined as:

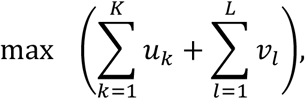

subject to:

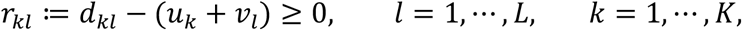

where for the dual variables *u*_*k*_ and *v*_*l*_, the reduced cost *r*_*kl*_ is defined as *d*_*kl*_ − (*u*_*k*_ + *v*_*l*_). The dual problem reformulates our optimization task to find an alternative reduction of the cost function with maximum sum and non-negative reduced costs. In summary, this algorithm constructs the auxiliary network (or equivalently a bipartite graph) and determines from an unassigned row *k* to an unassigned column *j* an alternative path of minimal total reduced cost and uses it to augment the solution^26^. In practice, despite time complexity *O*(*L*^3^), the Jonker-Volgenant algorithm is significantly faster than the majority of available algorithms for solving the assignment problem. By default, CytoSPACE calls the lapjv solver from the *lapjv* software package (version 1.3.14) in Python 3, which makes use of AVX2 intrinsics for speed (https://ms609.github.io/TreeDist/reference/LAPJV.html)^26^. With this solver, CytoSPACE runs in approximately 5 minutes on a single core using a 2.4 GHz Intel Core i9 chip for a standard 10x Visium sample with an estimated average of 5 cells per spot.

We provide an alternate solver based on the cost scaling push-relabel method^29^ using the Google OR-Tools software package in Python 3. This solver is an integer programming approximation method in which exact costs are converted to integers with some loss of numerical precision and which runs with time complexity *O*(*L*^2^ log(*LC*)), where *C* denotes the largest magnitude of an edge cost. In practice, this solver is approximately as fast as the Jonker-Volgenant based solver detailed above. However, for very large numbers of cells to be mapped, it can offer faster runtimes. Furthermore, it is supported more broadly across operating systems, so we recommend this solver for users working on systems which do not support AVX2 intrinsics as required by the lapjv solver. For users who wish to obtain the exact results of lapjv on operating systems that do not support the *lapjv* package, an equivalent but considerably slower solver implementing the Jonker-Volgenant algorithm is provided via the *lap* package (version 0.4.0), which has broad compatibility.

### Estimating cell type fractions

To overcome variability in cell type fractional abundance between a given ST sample and a reference scRNA-seq dataset, the first step of CytoSPACE requires estimating cell type fractions in the ST sample (**Fig. 1a**). Of note, only global estimates for the entire ST array are required and these may be obtained by combining spot-level fractions by cell type. Many deconvolution methods have been proposed to determine cell type composition from ST spots^14,20,27,28^, and any such method can be deployed for this purpose. In this paper, we used Spatial Seurat^14^ from Seurat version 3.2.3 for our primary analyses and show that correlations between estimated and true fractions of distinct cell types are high in simulated data (**Supplementary Fig. 3a**). After loading raw count matrices, we performed SCTransform() and RunPCA() with default parameters, followed by FindTransferAnchors() in which the preprocessed scRNA-seq and ST data served as the reference and query respectively. We then obtained spot-level predictions by TransferData() and obtained global predictions by summing prediction scores per cell type across all spots and scaling the sum of cell type scores to one.

In addition to Spatial Seurat, we tested the performance of RCTD^18^ for estimating global cell type fractions as input to CytoSPACE (**Supplementary Fig. 6**). RCTD version 2.0.0 (package spacexr in R) was employed with doublet_mode = ‘full’ and otherwise default parameters to obtain cell type fraction estimates per spot, followed by summing spot normalized result weights per cell type across all spots and scaling the sum to one.

### Estimating the number of cells per spot

The number of detectably expressed genes per cell (‘gene counts’) tightly corresponds to total captured mRNA content, as measured by the sum of unique molecular identifiers (UMIs) per cell^39^. As gene counts are routinely used as a proxy for doublets or multiplets in scRNA-seq experiments, we hypothesized that the sum of UMIs per ST spot may reasonably approximate the number of cells per spot, as required for the second step of CytoSPACE (**Fig. 1a**). To test this hypothesis while blunting the effect of outliers and technical variation, we first normalized UMIs to counts per million per spot and then performed log_2_ adjustment. We then estimated the number of cells per ST spot by fitting a linear function through two points: for the first point, we assumed that the minimum number of cells per spot is one and that this minimum in cell number corresponds to the minimum sum of UMIs in log_2_ space. For the second point, we assumed that the mean number of cells per spot corresponds to the mean sum of UMIs in log_2_ space and set this value according to user input. For 10x Visium samples in which spots generally contain 1-10+ cells per spot, we employed a mean of 5 cells per spot throughout this work. For legacy ST samples with larger spot dimensions, we selected a mean of 20 cells per spot. The number of cells for every spot was calculated from this fitted function. In support of our hypothesis, for simulated ST datasets, we found that the Pearson correlation between the estimated and real number of cells ranged between 0.80 and 0.93, depending on the dataset and spot resolution evaluated, validating our approach (**Supplementary Fig. 3b**).

### Harmonizing the number of cells per cell type

The third step of CytoSPACE equalizes the number of cells per cell type between the query scRNA-seq dataset and the target ST dataset (**Fig. 1a**). This is accomplished by sampling the former to match the predicted quantities in the latter using one of the following methods:

#### Duplication

Let *num*_*sc,k*_ and *num*_*ST, k*_ denote the real and estimated number of cells per cell type *k* in scRNA-seq and ST data, respectively. For cell type *k*, if *num*_*sc,k*_ < *num*_*ST,k*_, CytoSPACE retains all available cells in the scRNA-seq data and, also, randomly samples *num*_*ST,k*_ − *num*_*sc,k*_ cells from the same *num*_*sc,k*_ cells. Otherwise, it randomly samples *num*_*ST,k*_ from the *num*_*sc,k*_ available cells with cell type label *k* in the scRNA-seq data. By default, CytoSPACE applies this method for real data to ensure all cells assigned are biologically appropriate.

#### Generation

Here, when *num*_*sc,k*_ < *num*_*ST,k*_, instead of duplicating cells, new cells of a specific type are generated with independent random gene expression levels by sampling each gene from the gene expression distribution of cells of the same type uniformly at random. We used this method for benchmarking simulations to avoid bias in measuring precision owing to the presence of duplicated cells (**Fig. 1b-d**; **Supplementary Figs. 4, 5**, and **6**).

### Simulation framework

To evaluate the accuracy and robustness of CytoSPACE (**Fig. 1b**), we simulated ST datasets with known single-cell composition using Slide-seq datasets of mouse cerebellum and hippocampus sections^30^. Let *Sl* be an *M* × *B* gene expression matrix of a Slide-seq puck with *M* genes and *B* beads. To create a higher gene coverage version of *Sl*, denoted *Sc*, we used scRNA-seq datasets of the same brain regions^31^ to replace *Sl* beads with single-cell transcriptomes. Following quality control, in which outlier cells with >1,500 genes were removed, we matched each bead in the Slide-seq datasets with the nearest cell of the same cell type in the scRNA-seq dataset by Pearson correlation. We did this separately for each mouse brain region. As single cells may be matched with more than one bead, to obtain unique single-cell transcriptomes, we permuted genes between cells of the same cell type. For each cell, we replaced 20% of its transcriptome, with genes randomly selected per cell, with that of another randomly selected cell of the same cell type such that the latter is not a duplicate of the former. For simplicity, we matched the number of beads present in the two tissues by randomly sampling beads from the hippocampus data down to the number present in the cerebellum data.

Having created an *Sc* matrix for each brain region, we next sought to generate ST datasets with defined spot resolution. For this purpose, we imposed an *m* × *n* spatial grid over the entire puck. In each grid spot *x*_*ij*_, *i* = 1, …, *n, j* = 1, …, *m*, we calculated the sum of raw counts 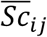 of the cells located within the grid-spot *x*_*ij*_. Since the spatial resolution of ST data varies depending on the technology used, we simulated ST datasets with an average of 5, 15, and 30 cells per spot.

Finally, in order to (1) leverage the scRNA-seq data underlying each *Sc* matrix as a query dataset and (2) emulate technical variation between platforms, we added noise to the scRNA-seq data in defined amounts. To this end, we selected a percentage of genes *p* to perturb, then randomly selected a corresponding subset of genes from each cell to which noise was added from the exponentiated Gaussian distribution 2^*N*(0,1)^. We considered noise perturbations for the following values of *p*: 5%, 10%, and 25%. Despite the addition of noise, UMAP plots of perturbed transcriptomes remained similar to the original data, implying maintenance of biologically-realistic data structure (**Supplementary Fig. 2**).

### Benchmarking analysis with simulated datasets

To benchmark CytoSPACE against Tangram, CellTrek, and two naïve distance metrics on simulated ST data, each approach was applied as follows:

#### CytoSPACE

For each ST resolution and scRNA-seq noise level, we estimated the fractional abundance of known cell types in the ST sample via Spatial Seurat, as described in “Estimating cell type fractions”. We then ran CytoSPACE with the “generated cells” option and with the lapjv solver implemented in Python (package *lapjv*, version 1.3.14).

#### Tangram

To ensure a fair comparison with CytoSPACE, we ran Tangram (version 1.0.2) with the same input cells mapped by CytoSPACE, including cells newly generated after resampling to match predicted cell type numbers. We also provided a normalized vector of CytoSPACE’s cell number per spot estimate as the density prior (density_prior argument). We trained Tangram on CPM-normalized scRNA-seq data in two ways: (1) using all available genes per cell and (2) using the top marker genes stratified by cell type. To identify marker genes using Seurat (version 4.1.0), we applied NormalizeData() with default parameters and FindAllMarkers() with only.pos = TRUE, min.pct = 0.1, and logfc.threshold = 0.25. The top 100 genes by average log_2_ fold change were then selected for each cell type.

#### CellTrek

Given that CellTrek heavily duplicates input cells (by default) and also filters input cells based on whether mutual-nearest neighbors are identified between cells and spots^25^, we provided CellTrek (version 0.0.0.9000) with all cells present in each simulated ST dataset (without the newly generated cells mapped by CytoSPACE and Tangram). After single cells were assigned to spatial coordinates, we selected the closest ST spot for each cell via Euclidean distance. As the CellTrek wrapper does not handle ST input without associated h5 and image files, we modified the code to accommodate ST datasets from other sources. CellTrek was run with default parameters, with the exception of (1) limiting the repel functionality (repel_r = 0.0001), as this parameter forces imputed spatial coordinates to arbitrarily deviate from their original predictions, and (2) setting spot_n to twice the mean number of cells per spot for each spatial resolution tested.

In addition to the above methods, we tested Euclidean distance and Pearson correlation. Here, each cell was assigned to the spot that either minimized distance or maximized correlation, respectively. All ground truth cells were evaluated without resampling and input datasets were CPM normalized and log_2_-adjusted prior to analysis.

#### Performance assessment

To determine the accuracy of single-cell mapping (**Fig. 1d, Supplementary Figs. 5 and 6**), we classified assigned locations that exactly matched ground truth spots as correct. Letting *TP*_*sc*_ denote the number of correct assignments, we defined single-cell precision (*Pr*_*sc*_) as

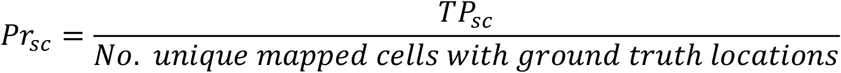

Of note, since generated cells (see “Harmonizing the number of cells per cell type”) did not have a corresponding ground truth location, they were excluded from this calculation. Separately, although CellTrek can assign the same cell ID *i* to multiple spots, any cell of ID *i* mapped to the correct spot at least once was considered correct. This was done without inflating the denominator or penalizing incorrect mappings for other cells with ID *i*.

### ST datasets for TME community analysis

Melanoma ST data generated by Thrane et al.^33^ were downloaded from https://www.spatialresearch.org/resources-published-datasets/doi-10-1158-0008-5472-can-18-0747/. Pre-processed spatial transcriptomics datasets of breast cancer (Visium fresh-frozen and FFPE) and colorectal cancer (fresh-frozen) specimens were downloaded from 10x Genomics (https://www.10xgenomics.com/spatial-transcriptomics/). Annotations of regions containing tumor cells were downloaded from 10x Genomics for the Visium FFPE breast cancer sample and shared by 10x Genomics upon request for the Visium fresh-frozen breast cancer sample analyzed in this work. A pre-processed Visium array of a fresh/frozen TNBC specimen (1160920F) was obtained from Wu et al.^35^ along with tumor boundaries.

### scRNA-seq tumor atlases

All analyzed tumor scRNA-seq data, which were downloaded as preprocessed count (UMI-based) or transcript (non-UMI-based) matrices, were selected and curated to clinically-match the ST specimens analyzed in this work (see “Molecular classification of breast cancer specimens”). Additionally, author-supplied annotations were used for all scRNA-seq reference datasets analyzed in Figure 2, with the following modifications. For the melanoma dataset generated by Tirosh et al.^34^, we excluded normal melanocytes and divided T cells into CD4 and CD8 subsets by the expression of *CD8A/CD8B* and *CD4/IL7R*, respectively, as previously described^4^. For the breast cancer dataset from Wu et al.^35^ and in the colorectal cancer dataset from Lee et al.^32^, the authors’ annotations were mapped to the major cell types analyzed in Figure 2. Of note, we excluded T cells that could not be confidently classified as CD8 or CD4 T cells and myeloid cells that could not be confidently classified as monocytes/macrophages or dendritic cells.

### Molecular classification of breast cancer specimens

When available, author annotations were used to determine estrogen receptor (ER) and human epidermal growth factor receptor 2 (HER2) enrichment status for each scRNA-seq and ST tissue breast cancer sample. For the FFPE breast cancer specimen from 10x Genomics without receptor status annotation, we examined the expression of *ESR1* (ER) and *ERBB2* (HER2) genes. We reclassified the FFPE breast cancer ST specimen as HER2+/ER-based on high expression of *ERBB2* without appreciable *ESR1* expression.

### Mapping of single-cell transcriptomes onto tumor ST samples

For the analyses in **Figure 2** and **Supplementary Figures 7–10**, CytoSPACE, Tangram, and CellTrek were applied as follows:

#### CytoSPACE

Cell type fractions were computed using Spatial Seurat (“Estimating cell type fractions”) and CytoSPACE was run with the “duplicated cells” option and the lapjv solver as implemented in the *lapjv* Python package on a single CPU core. For all Visium samples, we set the mean number of cells per spot to 5, while for legacy ST samples (melanoma ST data), we set this parameter to 20.

#### Tangram

As input, we analyzed the same single-cell transcriptomes mapped by CytoSPACE, including duplicates, along with a density prior (density_prior argument) determined by the number of cells per spot estimated by CytoSPACE. Since Tangram performed best with all genes when used for simulated ST datasets (**Fig. 1d, Supplementary Figs. 5 and 6**), we ran Tangram (version 1.0.2) on CPM-normalized scRNA-seq data with 24 CPU cores on all available genes. Other parameters were set to default.

#### CellTrek

Given CellTrek’s internal filtering mechanism (see “Benchmarking analysis with simulated datasets”), we provided all cells in the corresponding scRNA-seq atlases as input (without duplication or down-sampling). For Visium samples, we ran CellTrek (version 0.0.0.9000) with default parameters with 24 CPU cores (reduction=’pca’, intp=T, intp_pnt=10000, intp_lin=F, nPCs=30, ntree=1000, dist_thresh=0.4, top_spot=10, spot_n=10, repel_r=5, repel_iter=10, keep_model=T) and then assigned cells from raw output coordinates to their nearest spot by Euclidean distance. For the legacy ST samples (melanoma), we modified the code to handle inputs without h5 and image files, as detailed above. To fit the larger spot resolution in the legacy ST datasets, we fixed spot_n to 40. Other parameters were the same as above.

### Running time analysis

To evaluate the efficiency of CytoSPACE in practice and benchmark against Tangram and CellTrek, we recorded running times for each method across all scRNA-seq tumor atlas/ST pairs tested (n = 4 pairs with Visium ST data, n = 2 pairs with lower resolution legacy ST data) (**Supplementary Fig. 8a**) with parameter details as described above. For CytoSPACE, we report running times for both exact (shortest augmenting path via the lapjv solver) and integer approximation solvers, and both with and without a Spatial Seurat preprocessing step for obtaining input cell type fractional abundances. Data loading and file writing steps were excluded from running times for all methods. Methods were tested on comparable though not identical systems, with CytoSPACE, Spatial Seurat preprocessing steps, and Tangram tested on a computing cluster providing Intel E5-2640v4 (2.4 GHz base and 3.4 GHz max frequencies, with an associated 128 GB RAM), Intel 5118 (2.3 GHz base and 3.2 GHz max frequencies, with an associated 191 GB RAM), and AMD 7502 (2.5 GHz base and 3.35 GHz max frequencies, with an associated 256 GB RAM) processors, and with CellTrek tested on a server with an Intel E5-2680v3 processor and an associated 230 GB RAM. With the exception of CytoSPACE, in which the core mapping function uses only a single core, all methods were provided with 24 cores.

### Validation of alternative solver

To verify that the integer approximation solver we provide as a fast alternative to the recommended exact solver (lapjv) yields comparable results, we measured the proportion of single cells mapped to the same location across the two solver methods. For each scRNA-seq tumor atlas/ST pair tested, we mapped the same single cells after preprocessing for duplication and downsampling to match the estimated cell type fractions in tissue via CytoSPACE with exact and integer approximation solvers, and we report the percentage of cells mapped to the same spot in each method (**Supplementary Fig. 8b**). For duplicated cells, no distinction was made between the copies.

### Spatial enrichment analysis

To determine whether single cells mapped to ST spots showed enrichment of known spatially-resolved gene expression programs, cells were first partitioned into two groups (‘close’ and ‘far’) based on their distance from cancer cells. For breast cancer ST samples, all of which were profiled by 10x Visium, we used tumor boundary annotations determined by a pathologist in order to group cells. For melanoma and CRC datasets, the mean Euclidean distance of each TME cell to the nearest five tumor cells (mapped by the respective alignment method) was determined. For the melanoma dataset, melanoma cells were considered as tumor cells, while in the CRC dataset, tumor epithelial cells were considered for the purpose of identifying tumor locations in tissue. For each TME cell type, the resulting distances were median-stratified into ‘close’ and ‘far’ groups. This was done for two main reasons. First, the CRC sample lacked tumor boundary annotations. Second, while melanoma datasets included such annotations, the low spatial resolution of the legacy ST platform prevented precise co-registration with spatial spots at the tumor/stroma interface. That said, spatial enrichment results were comparable regardless of which distance method we employed for the melanoma datasets (data not shown).

To quantify spatial enrichment, we used pre-ranked gene set enrichment analysis (GSEA) implemented in fgsea (v1.14.0). As input, all spatially-mapped single-cell transcriptomes were loaded by cell type into Seurat v4.1.0 (min.cells = 5) and normalized with NormalizeData(). For each method and cell type, we then generated a gene list ranked by log_2_ fold-change for the identity classes “near” and “far” using FoldChange(). If fewer than 10 cells of a cell type were assigned to spots within one partition by at least one method, we excluded that cell type from the enrichment analysis. As CytoSPACE and Tangram were each run with the same scRNA-seq input, prior to running Seurat and fgsea, we performed random sampling of cells mapped by CellTrek in order to match the number of cells per cell type mapped by CytoSPACE and Tangram. This was done as described in “Harmonizing the number of cells per cell type - Duplication” in order to ensure a fair comparison among methods. Gene sets for T cell exhaustion and CE9/CE10-associated cell states were derived by Zheng et al.^37^ and Luca et al.^4^, respectively.

### Spatially-resolved macrophage states

To evaluate the spatial localization of *TREM2*^+^ and *FOLR2*^+^ macrophages^6^ (**Supplementary Fig. 10**), single-cell transcriptomes annotated as “Macrophages/Monocytes” were mapped to ST spots as described above (“Mapping of single-cell transcriptomes onto tumor ST samples”) and ordered based on their spatial distance (Euclidean) from tumor cells. All cells were processed with Seurat as described in “Spatial enrichment analysis”. To calculate distance, we used the same metric described for melanoma and CRC datasets (“Spatial enrichment analysis”). For cells mapped within tumor boundaries annotated by a pathologist (breast cancer datasets), distances were set to zero. We then divided cells into ‘near’ (distance = 0) and ‘far’ (distance > 0) groups and calculated the log_2_ fold change of each gene using FoldChange() in Seurat (**Supplementary Fig. 10b**).

### Statistics

All statistical tests were two-sided unless stated otherwise. The Wilcoxon test was used to assess statistical differences between two groups. Linear concordance was determined by Pearson (*r*) correlation or Spearman correlation (*ρ*), and a two-sided *t* test was used to assess whether the result was significantly non-zero. All statistical analyses were performed using R v3.5.1 and 4.0.2+, Python 3.8, MATLAB_R2019a, and Prism 9+ (Graphpad Software, La Jolla, CA).

### Code availability

CytoSPACE v1.0 was coded in Python and used to generate the results in this work. It is freely available, along with documentation, vignettes, and helper R scripts for creating CytoSPACE inputs and for estimating cell type fractions with Seurat, at https://github.com/digitalcytometry/cytospace.

## Notes

https://github.com/digitalcytometry/cytospace

### Competing Interest Statement

The authors have declared no competing interest.

## References

1. Keren, L., et al. A Structured Tumor-Immune Microenvironment in Triple Negative Breast Cancer Revealed by Multiplexed Ion Beam Imaging. Cell 174, 1373–1387 e1319 (2018).

2. Schürch, C.M., et al. Coordinated Cellular Neighborhoods Orchestrate Antitumoral Immunity at the Colorectal Cancer Invasive Front. Cell 182, 1–19 (2020).

3. Jackson, H.W., et al. The single-cell pathology landscape of breast cancer. Nature 578, 615–620 (2020).

4. Luca, B.A., et al. Atlas of clinically distinct cell states and ecosystems across human solid tumors. Cell 184, 5482-5496.e5428 (2021).

5. Grünwald, B.T., et al. Spatially confined sub-tumor microenvironments in pancreatic cancer. Cell 184, 5577-5592.e5518 (2021).

6. Nalio Ramos, R., et al. Tissue-resident FOLR2+ macrophages associate with CD8+ T cell infiltration in human breast cancer. Cell 185, 1189-1207.e1125 (2022).

7. Hu, J., et al. Statistical and machine learning methods for spatially resolved transcriptomics with histology. Comput Struct Biotechnol J 19, 3829–3841 (2021).

8. Edsgard, D., Johnsson, P. & Sandberg, R. Identification of spatial expression trends in single-cell gene expression data. Nat Methods 15, 339–342 (2018).

9. Halpern, K.B., et al. Paired-cell sequencing enables spatial gene expression mapping of liver endothelial cells. Nat Biotechnol 36, 962–970 (2018).

10. Halpern, K.B., et al. Single-cell spatial reconstruction reveals global division of labour in the mammalian liver. Nature 542, 352–356 (2017).

11. Hie, B., Bryson, B. & Berger, B. Efficient integration of heterogeneous single-cell transcriptomes using Scanorama. Nat Biotechnol 37, 685–691 (2019).

12. Moncada, R., et al. Integrating microarray-based spatial transcriptomics and single-cell RNA-seq reveals tissue architecture in pancreatic ductal adenocarcinomas. Nat Biotechnol 38, 333–342 (2020).

13. Nitzan, M., Karaiskos, N., Friedman, N. & Rajewsky, N. Gene expression cartography. Nature 576, 132–137 (2019).

14. Stuart, T., et al. Comprehensive Integration of Single-Cell Data. Cell 177, 1888–1902 e1821 (2019).

15. Stuart, T. & Satija, R. Integrative single-cell analysis. Nat Rev Genet 20, 257–272 (2019).

16. Sun, S., Zhu, J. & Zhou, X. Statistical analysis of spatial expression patterns for spatially resolved transcriptomic studies. Nat Methods 17, 193–200 (2020).

17. Zhu, Q., Shah, S., Dries, R., Cai, L. & Yuan, G.C. Identification of spatially associated subpopulations by combining scRNAseq and sequential fluorescence in situ hybridization data. Nat Biotechnol (2018).

18. Cable, D.M., et al. Robust decomposition of cell type mixtures in spatial transcriptomics. Nature Biotechnology 40, 517–526 (2022).

19. Dong, R. & Yuan, G.C. SpatialDWLS: accurate deconvolution of spatial transcriptomic data. Genome Biol 22, 145 (2021).

20. Elosua-Bayes, M., Nieto, P., Mereu, E., Gut, I. & Heyn, H. SPOTlight: seeded NMF regression to deconvolute spatial transcriptomics spots with single-cell transcriptomes. Nucleic Acids Res 49, e50 (2021).

21. Svensson, V., Teichmann, S.A. & Stegle, O. SpatialDE: identification of spatially variable genes. Nat Methods 15, 343–346 (2018).

22. Lohoff, T., et al. Integration of spatial and single-cell transcriptomic data elucidates mouse organogenesis. Nature Biotechnology (2021).

23. Dries, R., et al. Giotto: a toolbox for integrative analysis and visualization of spatial expression data. Genome Biology 22, 1–31 (2021).

24. Biancalani, T., et al. Deep learning and alignment of spatially resolved single-cell transcriptomes with Tangram. Nature Methods 18, 1352–1362 (2021).

25. Wei, R., et al. Spatial charting of single-cell transcriptomes in tissues. Nature Biotechnology (2022). https://doi.org/10.1038/s41587-022-01233-1

26. Jonker, R. & Volgenant, A. A Shortest Augmenting Path Algorithm for Dense and Sparse Linear Assignment Problems. Computing 38, 325–340 (1987).

27. Kleshchevnikov, V., et al. Cell2location maps fine-grained cell types in spatial transcriptomics. Nature Biotechnology (2022). https://doi.org/10.1038/s41587-021-01139-4

28. Newman, A.M., et al. Determining cell type abundance and expression from bulk tissues with digital cytometry. Nat Biotechnol 37, 773–782 (2019).

29. Goldberg, A.V. & Kennedy, R. An efficient cost scaling algorithm for the assignment problem. Math. Program. 71, 153–177 (1995).

30. Rodriques, S.G., et al. Slide-seq: A scalable technology for measuring genome-wide expression at high spatial resolution. Science 363, 1463–1467 (2019).

31. Saunders, A., et al. Molecular Diversity and Specializations among the Cells of the Adult Mouse Brain. Cell 174, 1015–1030 e1016 (2018).

32. Lee, H.-O., et al. Lineage-dependent gene expression programs influence the immune landscape of colorectal cancer. Nature Genetics 52, 594–603 (2020).

33. Thrane, K., Eriksson, H., Maaskola, J., Hansson, J. & Lundeberg, J. Spatially Resolved Transcriptomics Enables Dissection of Genetic Heterogeneity in Stage III Cutaneous Malignant Melanoma. Cancer Research 78, 5970–5979 (2018).

34. Tirosh, I., et al. Dissecting the multicellular ecosystem of metastatic melanoma by single-cell RNA-seq. Science 352, 189–196 (2016).

35. Wu, S.Z., et al. A single-cell and spatially resolved atlas of human breast cancers. Nature Genetics 53, 1334–1347 (2021).

36. Wherry, E.J. & Kurachi, M. Molecular and cellular insights into T cell exhaustion. Nature Reviews Immunology 15, 486–499 (2015).

37. Zheng, C., et al. Landscape of Infiltrating T Cells in Liver Cancer Revealed by Single-Cell Sequencing. Cell 169, 1342-1356.e1316 (2017).

38. Tyler, S.R., Bunyavanich, S. & Schadt, E.E. PMD Uncovers Widespread Cell-State Erasure by scRNAseq Batch Correction Methods. bioRxiv, 2021.2011.2015.468733 (2021).

39. Gulati, G.S., et al. Single-cell transcriptional diversity is a hallmark of developmental potential. Science 367, 405–411 (2020).

