## Supplementary Figures for "Robust alignment of single-cell and spatial transcriptomes with CytoSPACE"

### Table of Contents:

**Supplementary Figures**


| | |
|---|---|
| Supplementary Figure 1 | CytoSPACE versus conventional methods for decoding the cellular composition of bulk ST data |
| Supplementary Figure 2 | Impact of controlled noise on scRNA-seq query data |
| Supplementary Figure 3 | Estimation of cell type fractions and the number of cells per spot in bulk ST data |
| Supplementary Figure 4 | CytoSPACE alignments for all cell types analyzed in simulated ST datasets |
| Supplementary Figure 5 | Extended benchmarking analysis with simulated ST data |
| Supplementary Figure 6 | Performance of CytoSPACE with RCTD |
| Supplementary Figure 7 | Single-cell RNA-seq data mapped onto ST profiles of diverse human tumor specimens |
| Supplementary Figure 8 | Running time analysis and solver comparison |
| Supplementary Figure 9 | Spatial enrichment of tumor-associated cell states across methods and datasets |
| Supplementary Figure 10 | Single-cell spatial analysis of *TREM2*$^+$ and *FOLR2*$^+$ macrophage states across methods and datasets |
| Supplementary Figure 11 | UMAP projections of scRNA-seq tumor atlases labeled by predicted spatial locations |

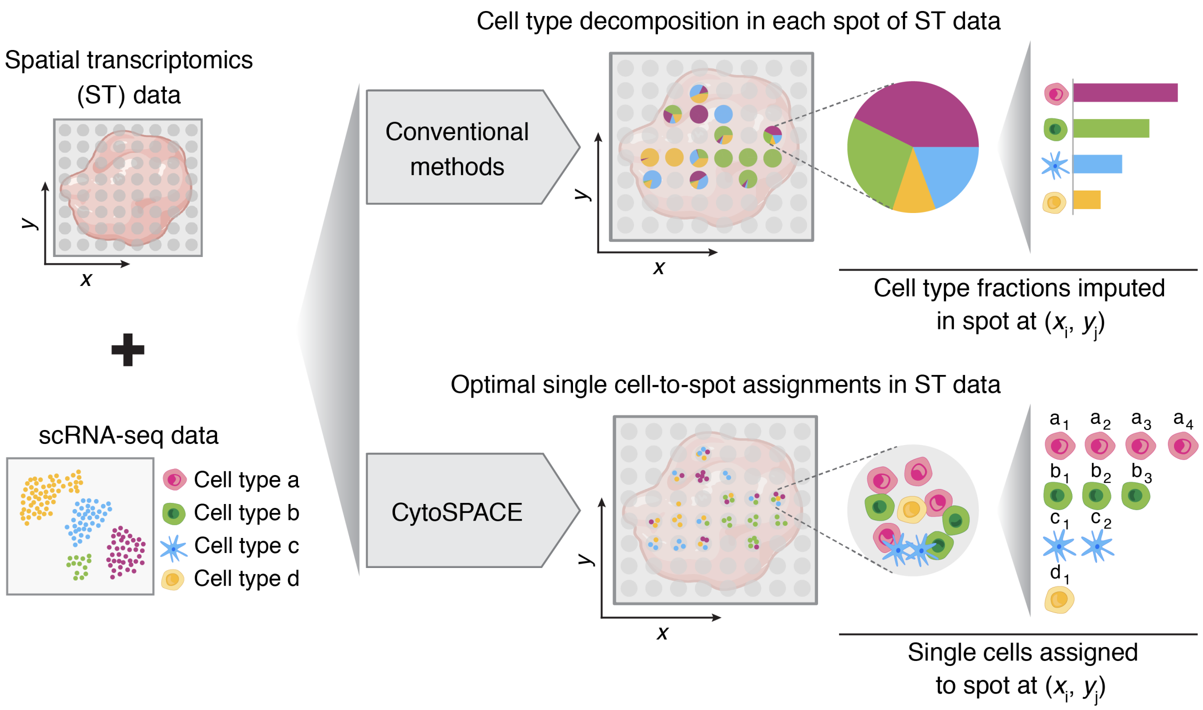


**Supplementary Figure 1: CytoSPACE versus conventional methods for decoding the cellular composition of bulk ST data.** Most methods for deconvolving bulk ST data estimate cell type fractions using single-cell reference profiles (top). In contrast, CytoSPACE efficiently assigns individual single-cell transcriptomes to ST coordinates (i.e., spots) using convex optimization to globally minimize a correlation-based cost function through a linear programming routine. This enables downstream analysis of cell type proportions and single-cell transcriptional heterogeneity in spatial dimensions (bottom). The labels $a_1, \ldots, d_1$ denote individual single cells of cell type $a, \ldots, d$, respectively, assigned to the featured spot.

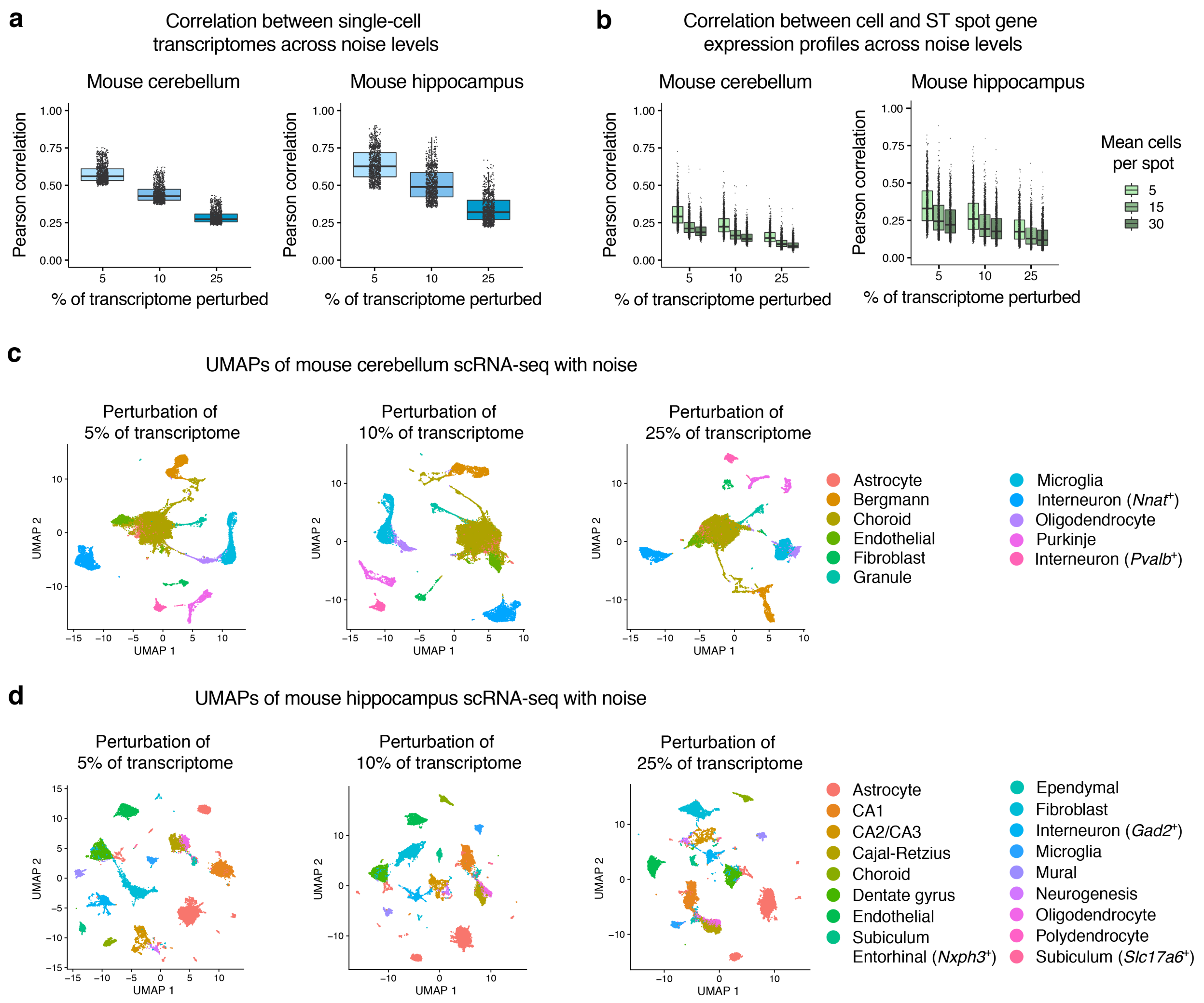


**Supplementary Figure 2: Impact of controlled noise on scRNA-seq query data. a**, Box plots showing the effect of adding noise to the scRNA-seq query datasets used in simulation experiments. In brief, single-cell expression profiles of mouse cerebellum and hippocampus were perturbed by adding noise sampled from an exponentiated normal distribution to randomly selected genes, comprising 5% to 25% of each cell's original transcriptome (x-axis, **Methods**). Concordance between the original and perturbed transcriptome in $\log_2$ space for 1,000 randomly sampled cells per scRNA-seq dataset (mouse cerebellum, left; mouse hippocampus, right), expressed as Pearson correlation coefficient (y-axis). The box center lines, box bounds, and whiskers indicate the medians, first and third quartiles and minimum and maximum values within 1.5× the interquartile range of the box limits, respectively. **b**, Same as a but showing Pearson correlation (y-axis) between 1,000 randomly selected single-cell transcriptomes after the addition of noise (x-axis) and their corresponding ground truth ST spot transcriptomes, for different mean spot resolutions (legend). Pearson correlation was determined in $\log_2$ space. **c-d,** UMAP embeddings of scRNA-seq query after the addition of noise for mouse cerebellum (c) and mouse hippocampus (d) datasets. Importantly, cell type clusters are maintained across the range of considered perturbations.

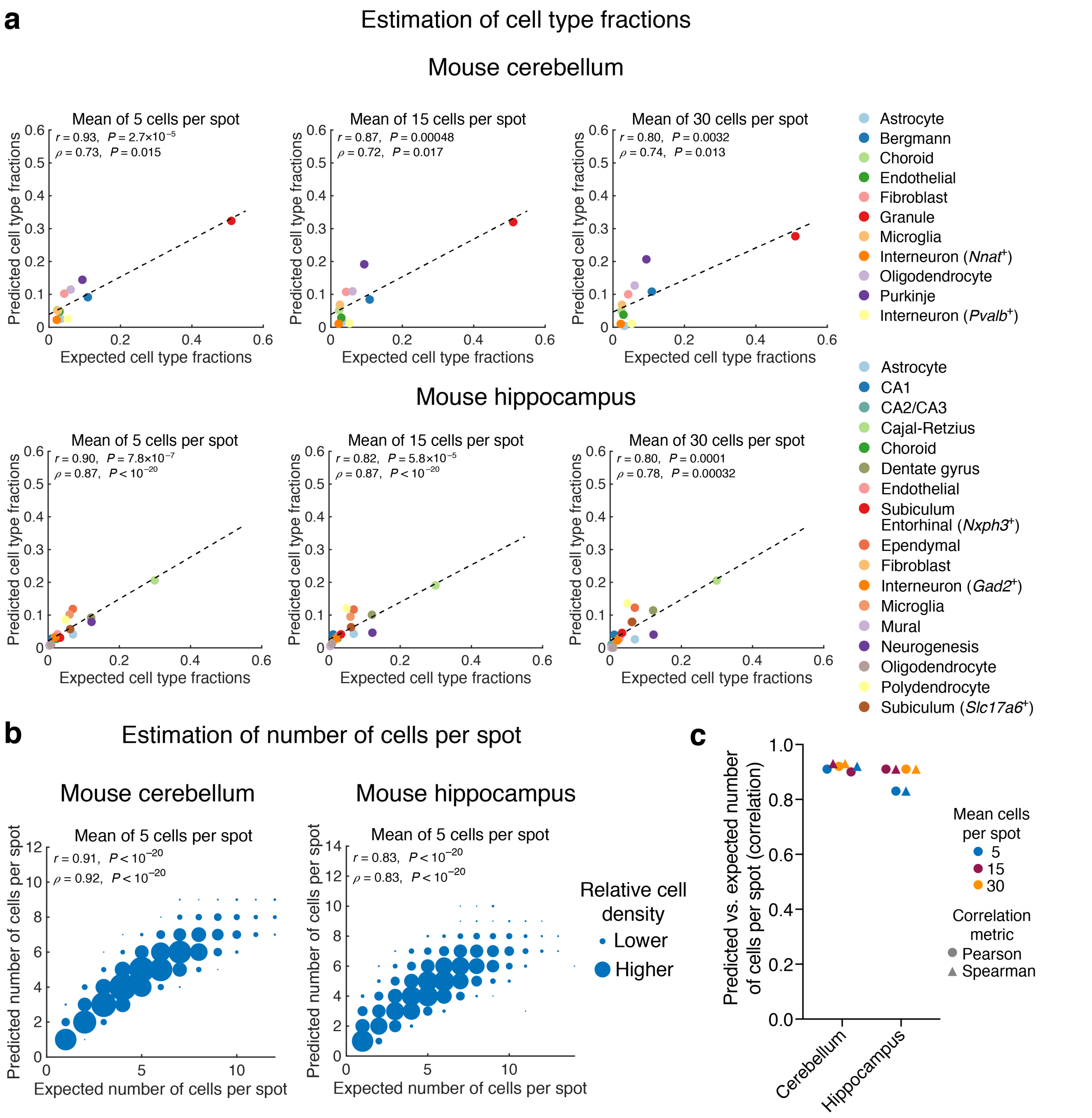


**Supplementary Figure 3. Estimation of cell type fractions and the number of cells per spot in bulk ST data. a**, Application of Spatial Seurat to infer cell type fractions in simulated ST datasets (**Methods**). Scatter plots show ground truth cell type fractions (x-axis) versus estimated fractions (y-axis) for simulated ST data of mouse cerebellum (top) and hippocampus (bottom) sections with different spot resolutions. Single-cell RNA sequencing data were first perturbed with the addition of noise to 5% of the transcriptome, as described in **Methods**. **b**, Scatter plot showing the number of cells per spot estimated by CytoSPACE in simulated ST datasets (y-axis; **Methods**) versus ground truth (x-axis) at a mean of 5 cells per spot for mouse cerebellum and hippocampus sections. Relative density is depicted by point size. Concordance and significance were assessed by Pearson *r* or Spearman ρ and a two-sided *t* test, respectively. **c**, Same as b but showing correlation coefficients (Pearson and Spearman) for all analyzed spot resolutions. All correlations are significant ($P < 10^{-20}$).

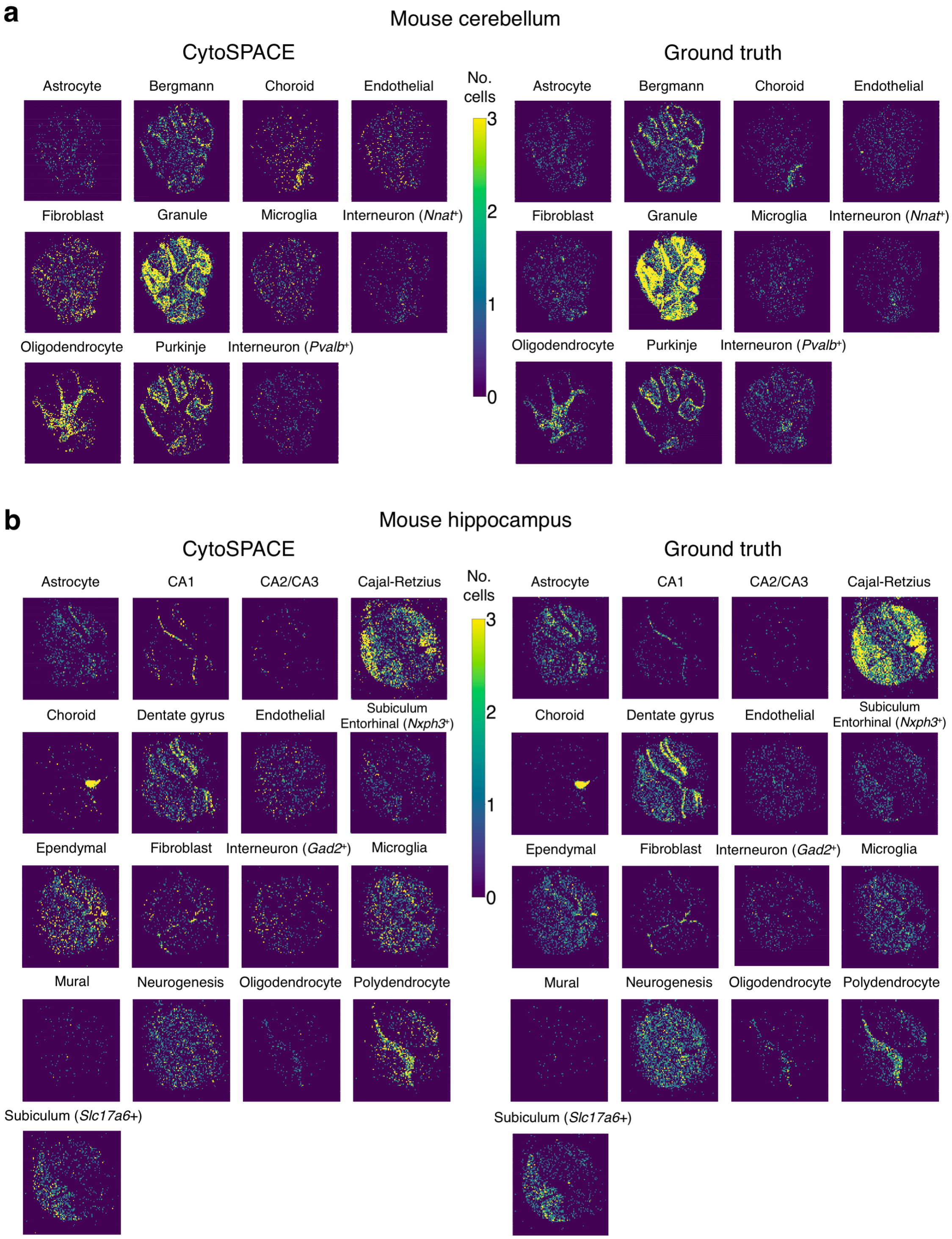


**Supplementary Figure 4: CytoSPACE alignments for all cell types analyzed in simulated ST datasets (related to Fig. 1c). a-b,** Same as Figure 1c but shown for all evaluated cell types mapped to mouse cerebellum (*n* = 11 cell types) (a) and hippocampus (*n* = 17 cell types) (b) ST datasets defined by simulation, with a mean of 5 cells per spot. CytoSPACE assignments, shown for single-cell transcriptomes with noise applied to 5% of the transcriptome (**Methods**), demonstrate strong concordance with ground truth.

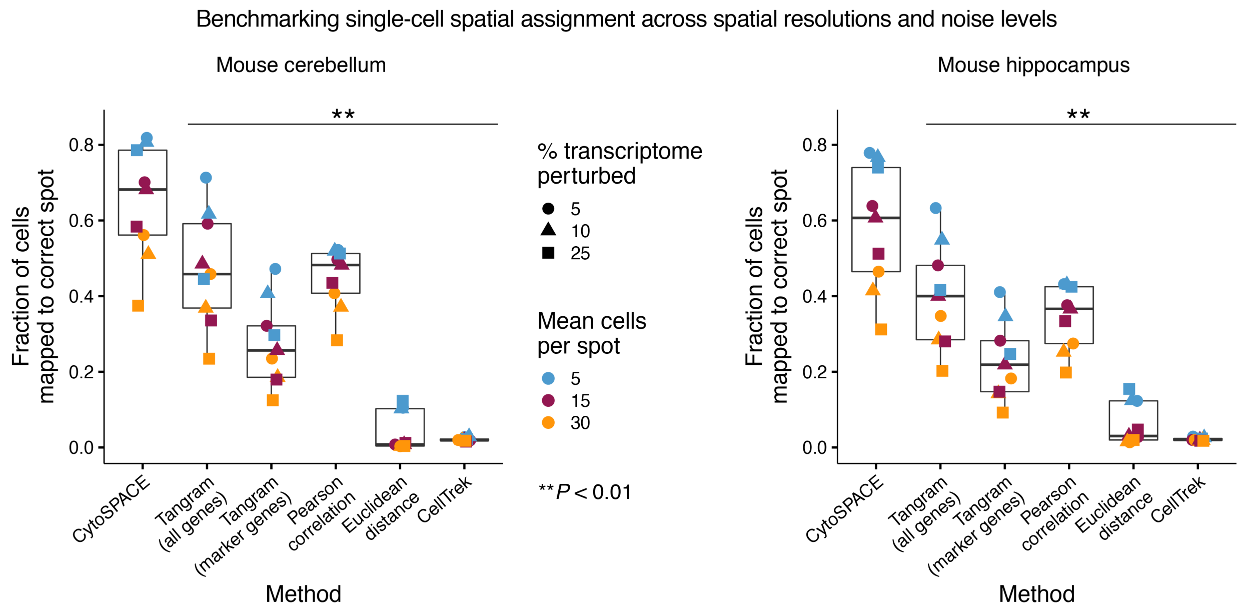


**Supplementary Figure 5: Extended benchmarking analysis on simulated ST data (related to Fig. 1d).** Box plots depicting the fraction of all single-cell transcriptomes assigned to the correct ST spot, shown for different spot resolutions (mean of 5, 15, and 30 cells per spot) and scRNA-seq noise levels (perturbations added to 5%, 10%, and 25% of the transcriptome) for each evaluated method. Statistical significance was determined using a two-sided paired Wilcoxon test (***P* < 0.01).

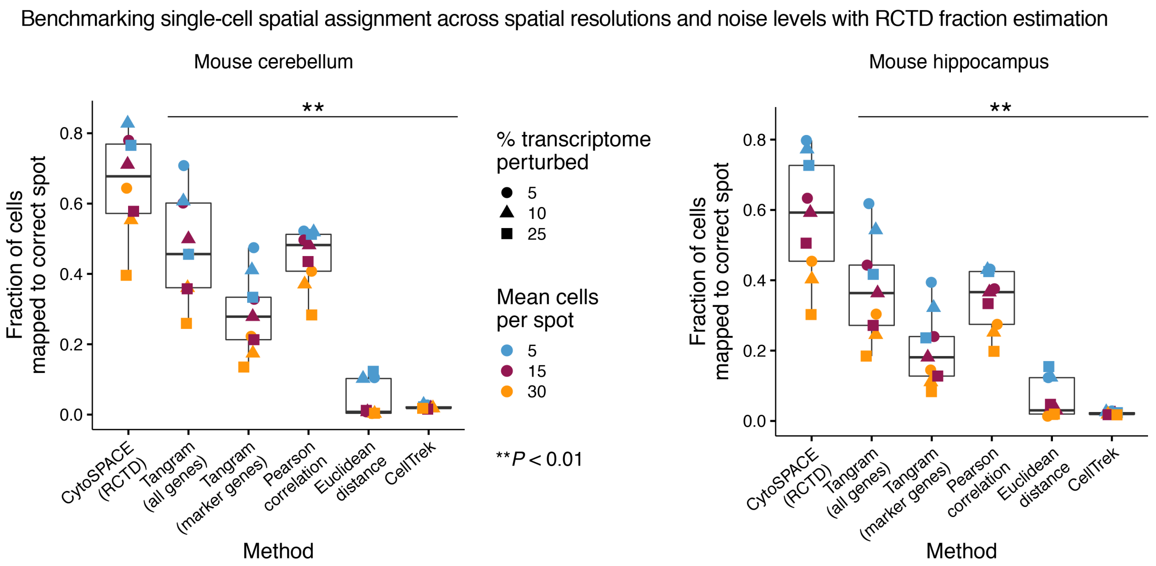


**Supplementary Figure 6: Performance of CytoSPACE with RCTD.** Same as Supplementary Figure 5 but showing the application of CytoSPACE with RCTD for cell type fraction estimation (rather than Spatial Seurat).

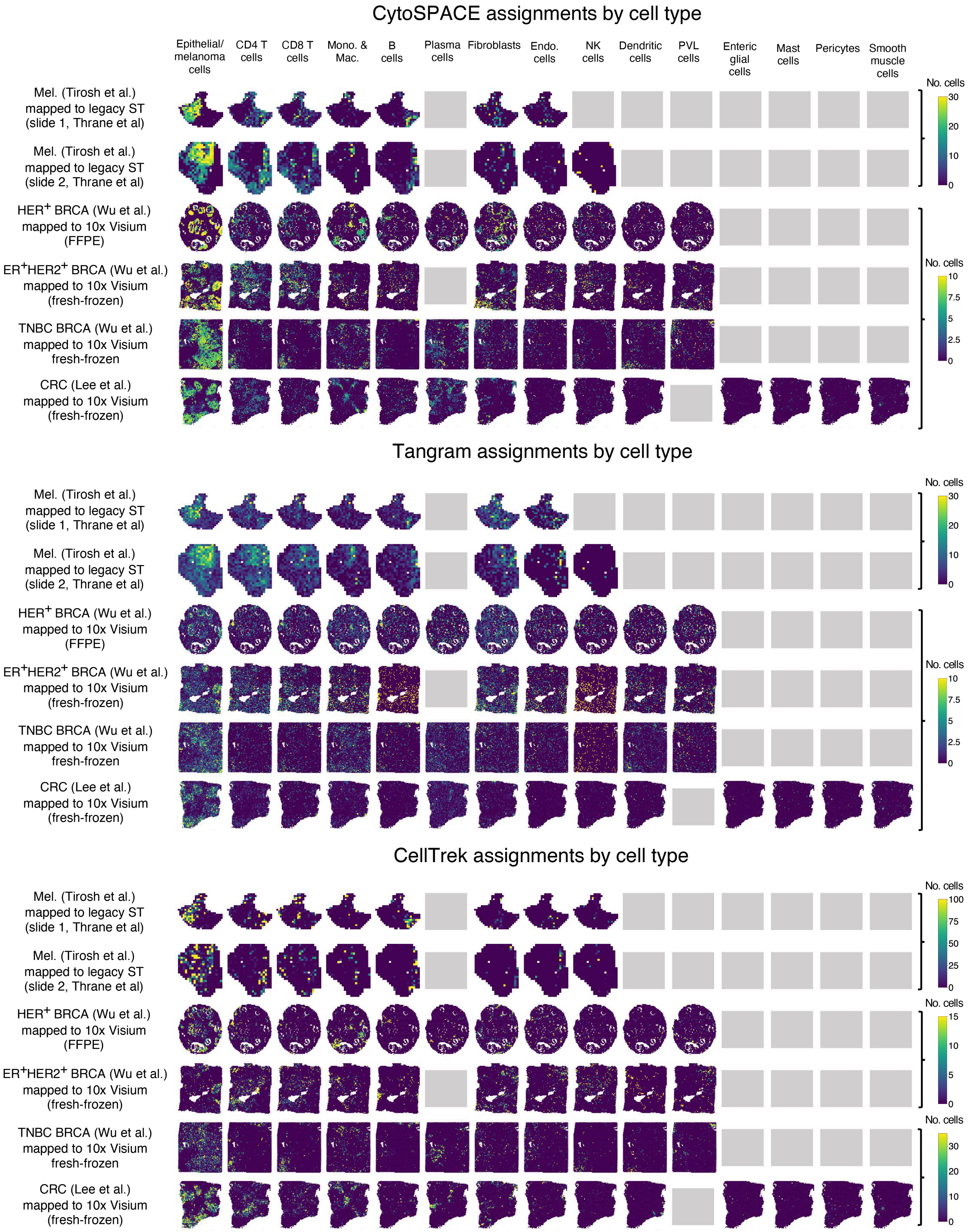


**Supplementary Figure 7: Single-cell RNA-seq data mapped onto ST profiles of diverse human tumor specimens.** Same as Figure 2a but showing all cell types analyzed for each scRNA-seq/ST dataset by CytoSPACE, Tangram, and CellTrek. Gray boxes denote cell types without author-supplied annotations in the corresponding scRNA-seq atlas (**Methods**).

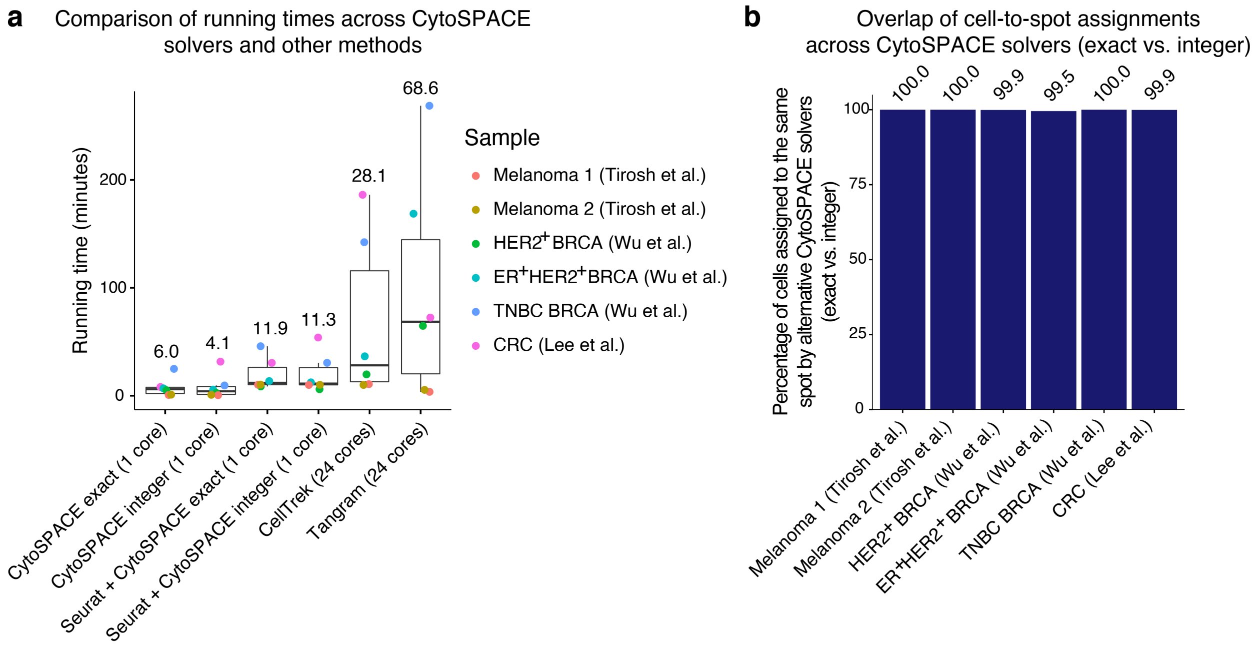


**Supplementary Figure 8: Running time analysis and solver comparison. a**, Comparison of running times across methods for representative scRNA-seq/ST datasets analyzed in this work. In all cases, the core CytoSPACE mapping function was run with a single CPU, whereas Tangram and CellTrek were each run with 24 CPU cores. For the fractional abundance inference step via Spatial Seurat, 24 CPU cores were provided. For all methods, data loading and file writing were excluded from reported running times. Median running times are reported above each group. For parameter details, see **Methods**. The box center lines, box bounds, and whiskers indicate the medians, first and third quartiles and minimum and maximum values within 1.5× the interquartile range of the box limits, respectively. **b**, Concordance between CytoSPACE solvers (exact shortest augmenting path vs. integer approximation cost scaling push-relabel methods) for single-cell spot assignment in selected datasets. In all cases tested, greater than 99% of cells were assigned to the same spot between solver methods.

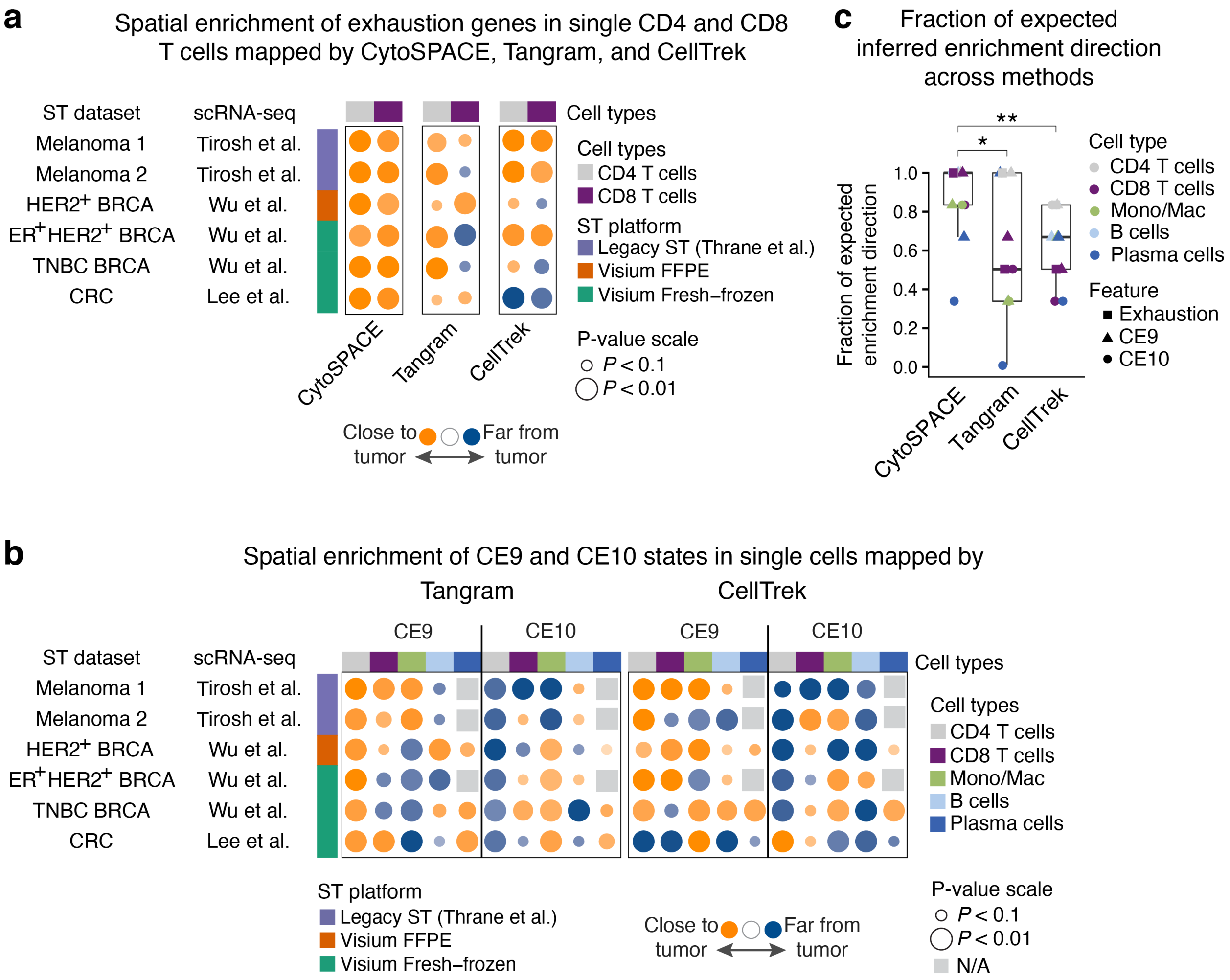


**Supplementary Figure 9: Spatial enrichment of tumor-associated cell states across methods and datasets. a**, Bubble plot showing the spatial enrichment of exhaustion genes in CD4 and CD8 T cell transcriptomes mapped onto ST spots by CytoSPACE, Tangram, and CellTrek (related to **Fig. 2d**). Bubbles denote normalized enrichment scores calculated by pre-ranked GSEA, as described in the caption of Figure 2b and in **Methods**. scRNA-seq/ST dataset pairs are ordered as in Figure 2f. **b**, Same as Figure 2f but showing performance for Tangram and CellTrek. Single-cell RNA-seq datasets without annotated plasma cells are indicated by gray boxes ("N/A"). **c**, Fraction of datasets per cell type for which the expected spatial enrichment direction was correctly inferred by CytoSPACE, Tangram, and CellTrek for each of the gene sets analyzed in this work ($n$ = 11 distinct gene sets with 12 data points per method, as canonical exhaustion genes were analyzed for CD4 and CD8 T cells). The box center lines, box bounds, and whiskers indicate the medians, first and third quartiles and minimum and maximum values within 1.5× the interquartile range of the box limits, respectively. Statistical significance was determined by a paired two-sided Wilcoxon test relative to CytoSPACE. *$P$ < 0.05, **$P$ < 0.01.

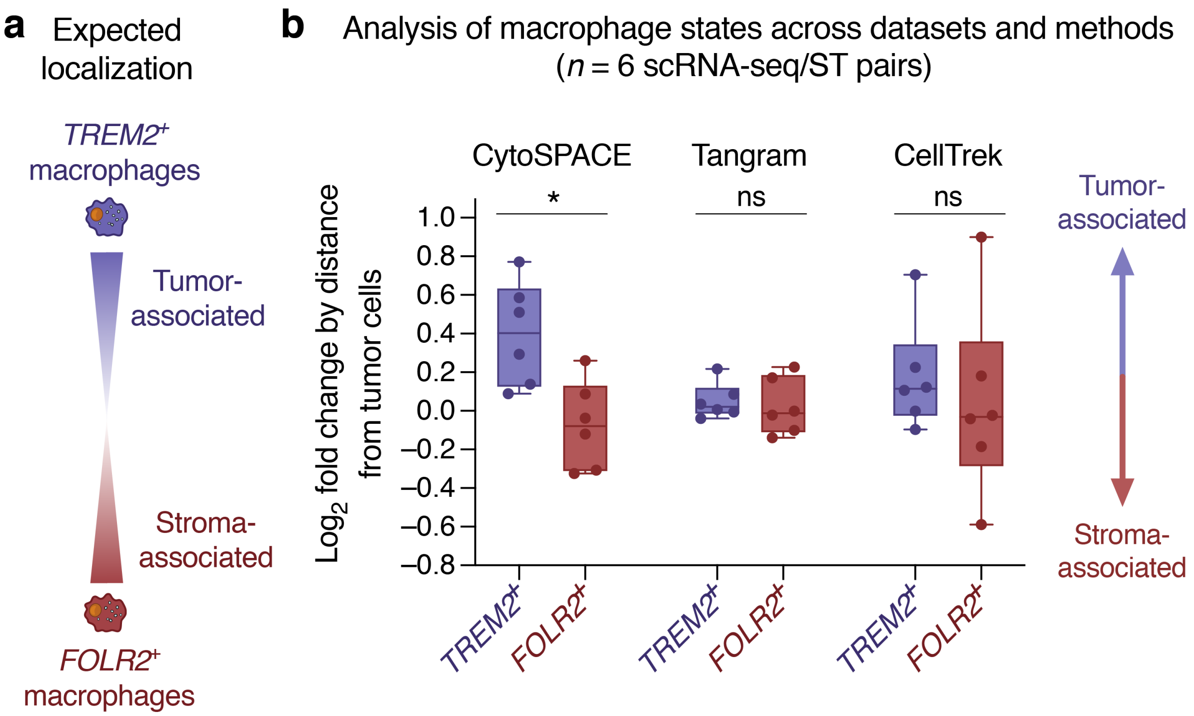


**Supplementary Figure 10: Single-cell spatial analysis of *TREM2*⁺ and *FOLR2*⁺ macrophage states across datasets and methods. a,** Expected spatial localization of *TREM2*⁺ and *FOLR2*⁺ macrophages in human tumors (Nalio Ramos et al.). **b**, Box plots comparing the $\log_2$ fold change of *TREM2* and *FOLR2* expression in single macrophage/monocyte transcriptomes grouped into 'near' (Euclidean distance to tumor = 0) and 'far' (Euclidean distance to tumor > 0) categories, as described in **Methods**. Each point represents an scRNA-seq/ST pair analyzed in Figure 2f. Single-cell mappings for each of the three methods are identical to Figure 2. The box center lines, box bounds, and whiskers denote the medians, first and third quartiles and minimum and maximum values, respectively. Two-group comparisons were performed using a two-sided paired Wilcoxon test (indicated by the horizontal line above each pair of *TREM2*⁺ and *FOLR2*⁺ boxes; $*P < 0.05$).

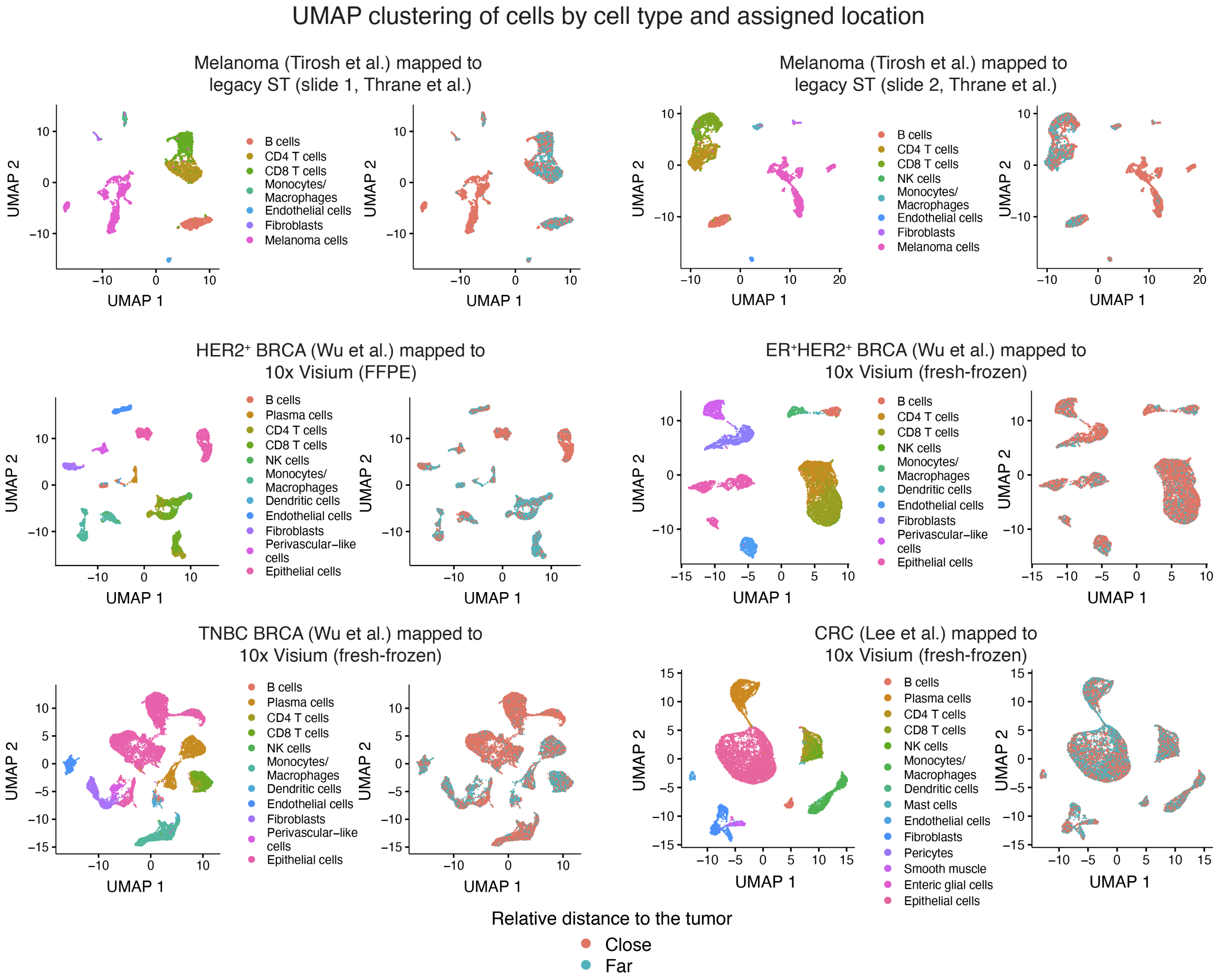


**Supplementary Figure 11: UMAP projections of scRNA-seq tumor atlases labeled by predicted spatial locations.** UMAP embeddings showing all single-cell transcriptomes mapped by CytoSPACE to ST samples analyzed in Figure 2. Cells are colored by lineage (left) and by relative distance to tumor cells (right), determined as described in **Methods**.
